# Intracameral Puncture Lowers Intraocular Pressure and Triggers an Immune Response in the Conventional Outflow Tract

**DOI:** 10.1101/2025.03.03.639599

**Authors:** Darren M. Schuman, Maria Fernanda Suarez, Aleksander O. Grimsrud, Megan Kuhn, Daniel R. Saban, W. Daniel Stamer, Katy C. Liu

## Abstract

**Purpose:** Intracameral injection is an effective delivery method for biomedical agents and therapeutics to conventional outflow tract tissues. However, the effect of intracameral injections on intraocular pressure and aqueous dynamics has not been well characterized, warranting further investigation.

**Methods:** Wild type 3-5-month-old C57BL/6 mice were subjected to intracameral puncture (ICP, without injection of any material). Following ICP, intraocular pressure (IOP), outflow facility, aqueous production, episcleral vessel diameter, and macrophage densities were measured.

**Results:** On day 1, IOP was significantly reduced by 30% (*p* < 0.0001; n=25) while outflow facility (*p* = 0.306; n=15) and aqueous production (*p* = 0.163; n=9) were unchanged. As well, Schlemm’s canal filtration area was unchanged, however distal vessels were dilated (*p* < 0.001) at day 1 post ICP. Correspondingly, macrophage density was significantly increased around episcleral vessels (*p* < 0.0005) at day 1. Macrophage densities in Schlemm’s canal and trabecular meshwork, while unchanged at day 1, were significantly increased by day 3 (*p* < 0.0001).

**Conclusions:** Coincident with significantly reduced IOP one day after ICP, there was an influx of macrophages into the distal portion of the conventional outflow tissues and a dilation of episcleral vessels, likely reducing distal outflow resistance. Our study demonstrates the importance of considering the drug delivery method to the eye due to its effects on the immune response and conventional outflow homeostasis.

## Introduction

Intracameral injections have been used as an effective delivery means to anterior ocular tissues such as the outflow tract (i.e., trabecular meshwork and Schlemm’s canal) and corneal endothelium.^1,2^ Drugs, viral vectors, beads and stem cells have been injected into the anterior chamber for research purposes in mice.^1,3–5^ In humans, intracameral injections have an extensive clinical history dating back 60 years with drugs such as antibiotics.^6,7^ Recently, the intracameral delivery method has been used to dispense the sustained-release Durysta^TM^ bimatoprost implant (Allergan, Dublin, IE) into the anterior chamber and localizing adjacent to targeted tissue, the trabecular meshwork.^8^ Yet, intracameral drug delivery is not without off-target effects. Injections of recombinant adenovirus into the mouse anterior chamber result in prolonged localization of macrophages to the iris and iridocorneal angle,^9^ which may alter study results and interpretation.

Macrophages are mononuclear phagocytes that reside in tissues throughout the body in homoeostasis and are recruited to tissues following injury or disease. Specifically, monocytes from the bone marrow are recruited and differentiate into monocyte-derived macrophages within the tissue. Macrophages reside in the conventional outflow tract tissues of the trabecular meshwork (TM), Schlemm’s canal (SC) and around distal vessels (DV) in homeostasis,^10,11^ and monocytes can be recruited to the TM after a stimulus such as laser trabeculoplasty.^11^ Moreover, laser trabeculoplasty affects intraocular pressure (IOP), which has been proposed mechanistically to be due to monocyte infiltration.^11^ Additionally, ocular inflammatory conditions or uveitis can cause IOP changes including abnormally low IOP or hypotony.^12^

While macrophages are key sensors following tissue injury,^13^ it is unknown whether intracameral injections result in macrophage recruitment to conventional outflow tissues within the anterior chamber. Furthermore, the effect of intracameral injections on IOP and aqueous humor dynamics has not been studied. Such physiologic changes are relevant to the interpretation of experimental data relating to the conventional outflow tract and inflammatory changes. Here, we evaluate IOP, aqueous humor dynamics and macrophage infiltration following intracameral needle puncture.

Our results show that intracameral needle puncture transiently reduces IOP and is associated with macrophage recruitment to and egress out of outflow tract tissues. The reduced IOP may be due to vasomotor changes to the episcleral venous system.

## Methods

### Animals

C57BL/6 wild type mice were purchased from The Jackson Laboratory (Bar Harbor, MA, USA). Mice were handled in accordance with Duke University Institutional Animal Care and Use Committee and in compliance with the Association for Research in Vision and Ophthalmology (ARVO) Statement for the Use of Animals in Ophthalmic and Vision Research. Mice were aged 3-5 months for all studies. Animals were fed a normal diet with unlimited access to food and water and subjected to a 12-hour light/dark cycle at 21°C.

### Intracameral Punctures

One eye per animal was punctured while the untreated contralateral eye served as control. Mice were anesthetized with isoflurane (2% v/v for induction; 1.0%–1.5% v/v for maintenance) delivered through a nose cone. The treated eye received one drop of 0.5% proparacaine (Sandoz) to the cornea, followed by 1 drop of a dilating agent (0.5% Tropicamide & 5% Phenylephrine; Duke compounding pharmacy). Once dilation of the eye was confirmed, a pulled glass needle connected to a 10 µL syringe (Hamilton, Franklin, MA, USA) on a micromanipulator was used to produce a full-thickness puncture through the cornea into the anterior chamber without hitting the lens (Fig. 1a). The needle remained in the anterior chamber for approximately 5 minutes followed by topical application of 0.5% erythromycin ophthalmic ointment (Akorn) and removal of the glass needle from the eye. The animal was then placed on a heating pad for recovery.

**Figure 1:**
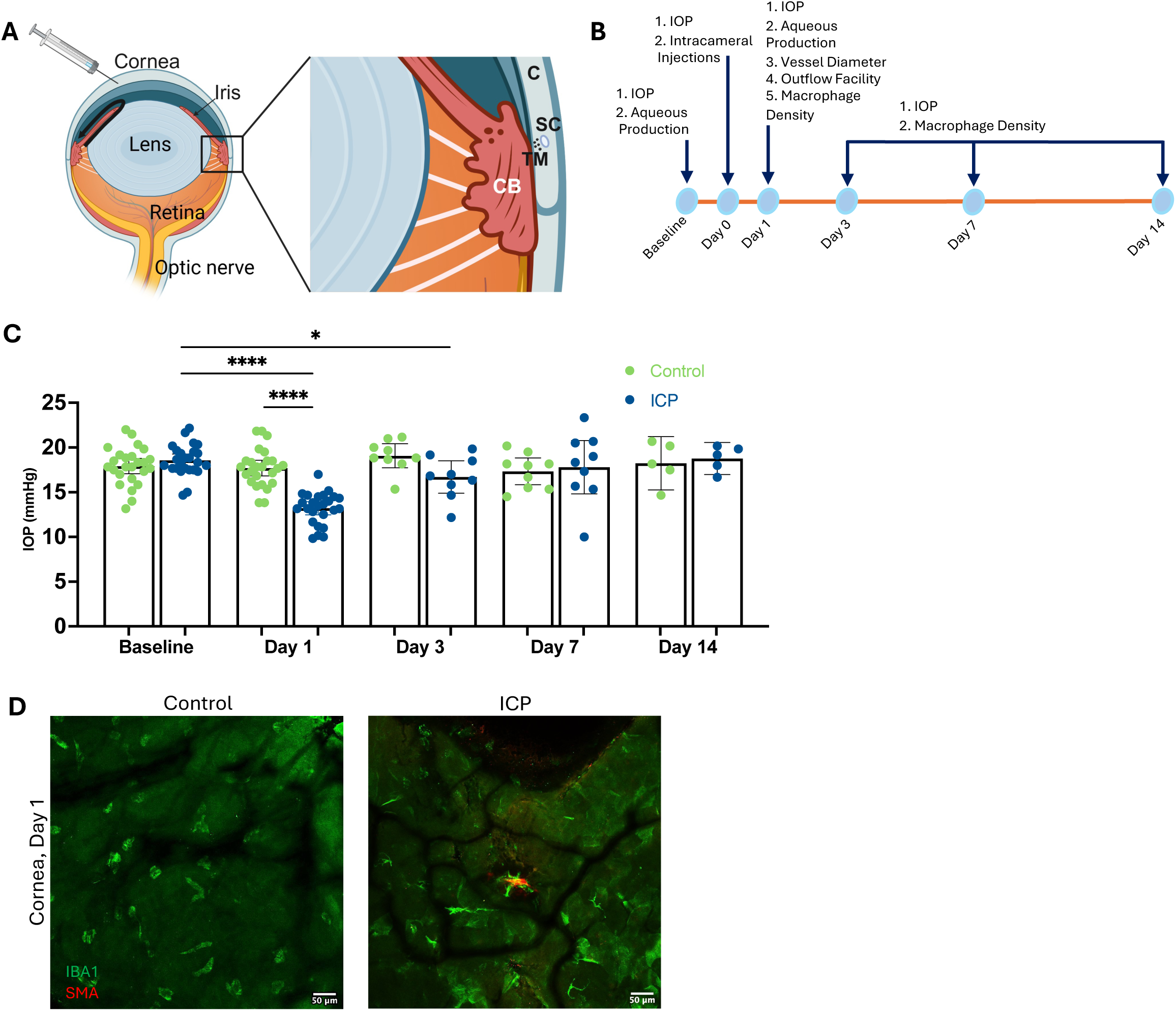
Intracameral corneal puncture lowers IOP at day 1. (A) Schematic of mouse eye with intracameral injection to ½ anterior chamber depth. Black arrow indicates aqueous humor flow (left) and magnified schematic of aqueous humor outflow tract (right). (B) Schematic of timeline used to measure IOP, aqueous production, episcleral vessel diameter, outflow facility and macrophage density. (C) IOPs in C57BL/6 mice, age 3-5 months, measured at baseline, days 1, 3, 7, and 14 following ICP with contralateral eyes as control. (D) Representative images of control (left) and ICP (right) cornea flat mounts immunostained with anti-Iba1 and anti-αSMA at day 1 post ICP (scale bar = 50 µm). ICP: intracameral puncture; C: cornea; SC: Schlemm’s canal; TM: trabecular meshwork; CB: ciliary body. * *p* < 0.04; **** *p* < 0.0001.

### IOP Measurements

Aerosolized isoflurane was delivered for IOP measurements as previously described.^14,15^ A Tonolab (iCare, Finland) rebound tonometer was used for all IOP measurements, taking no longer than 5 minutes after anesthesia induction. IOP for each eye was measured 6 times and averaged in data analysis. IOP was measured in the mornings for consistency.

### Aqueous Production Measurements

Aqueous humor production was measured with a Fluorotron Master for mice (Ocumetrics, Mountain View, CA) similar to previously described techniques.^16^ 10 µL of 2% fluorescein (Duke compounding pharmacy) was topically applied to conscious mice followed by a 1 hour wash out period. Mice were then anesthetized with ketamine (100 mg/kg)/xylazine (10 mg/kg). A second dose of ketamine/xylazine was delivered after 45 minutes. Fluorophotometry measurements were taken at least 4 times every 15 minutes on a 37.4°C heated platform. Anterior chamber volume of 3 µL was used as a standard to calculate aqueous production.^17^

### Outflow Facility Measurements

As described by Sherwood *et al.*, iPerfusion was used to measure outflow facility.^18^ Mice were euthanized 1 day following ICP. The eyes were then carefully enucleated and glued to a small platform in the perfusion eye bath. The baths were filled with prewarmed, degassed PBS with added glucose (DBG) and kept at a constant temperature. The eyes were cannulated with pulled, sharpened glass needles that were filled with prewarmed, degassed DBG and attached to the perfusion system. After cannulation, the eyes underwent a 30-minute acclimation at 12 mmHg then went through a series of pressure steps to measure facility. The steps started at 5 mmHg, increased 1.5 mmHg each step until reaching 17 mmHg, before decreasing to the final step at 8mmHg. iPerfusion software was used to analyze the perfusion data and generate a facility for each eye.

### Tissue Harvesting and Histology

Animals were sacrificed and eyes carefully enucleated. Eyes were fixed in 4% paraformaldehyde, dissected, and immunostained as anterior cup whole mounts. Samples were placed in a blocking solution (5% goat serum, 3% bovine serum albumin, 1% Triton-X, and 0.1 Tween-20) for 1 hour at room temperature followed by incubation in primary antibody solutions (1:50, Armenian hamster anti CD31, clone 2H8, Developmental Studies Hybridoma Bank, University of Iowa; 1:200, rabbit anti IBA1, Fujifilm, #019-19741, Osaka, Japan). After an overnight incubation at 4°C, samples were washed with 0.2% Tween in PBS. Samples were then incubated in secondary antibody solution (1:250, goat anti-Armenian hamster, #127-605-160; goat anti-rabbit, #112-545-167, Jackson ImmunoResearch Laboratories, Carlsbad, CA, USA) and 1:200 conjugated Cy3 anti-smooth muscle actin (Millipore, C6198). After overnight incubation at 4°C, samples were washed and mounted using Immuno-Mount^TM^ (ThermoFisher Scientific). For ciliary body characterization, approximately ¼ of an anterior cup was used for agarose sectioning after immunostaining. A vibratome was used to create 50 µm cross sections from 5% agarose molds. Tissue sections were placed on glass slides for imaging using a Nikon Eclipse Ti2 inverted confocal microscope (Nikon, Melville, NY, USA). Under 40x magnification, 5-6 images of the outflow tract and ciliary body were collected as nonconsecutive z-stacks. For distal vessel dimensional analysis, samples were reimaged as tiled z-stacks under 20x magnification.

### Macrophage and Vessel Diameter Quantification

IBA1^+^ macrophages were quantified in the ciliary body (CB) and TM, SC, and DV of the conventional outflow tract using ImageJ. Macrophages colocalizing with SMA were counted in TM, and macrophages adjacent to the CD31^+^ SC were counted in SC. DV macrophages adjacent to CD31^+^ episcleral vessels were counted. Ciliary body macrophages within ciliary epithelium were quantified. A freehand tracer tool was used to define the borders of SC and DV for area measurements. To measure episcleral vessel diameters, a 5×5 grid was overlaid in ImageJ and used as reference points to measure the diameters of episcleral vessels as previously described.^14^ Arterioles were identified as CD31^+^ with striated SMA staining, venules were CD31^+^ with non-striated SMA staining, and capillaries were smaller CD31^+^ connecting vessels.

### Statistical Analysis

Statistical analyses were performed using GraphPad Prism v.8 (GraphPad Software, La Jolla, CA). Student t-tests and Welch’s tests were used to compare averages of data sets with p < 0.05 as statistically significant. Data is presented as mean ± SEM.

## Results

### Intracameral Puncture Reduces IOP at Day 1

IOP was measured following ICP (day 0) on days 1 (24 hours), 3, 7 and 14 (Fig 1b). On day 1, IOP was reduced 30% after ICP compared to baseline (18.56 ± 0.35 mmHg to 13.21 ± 0.35 mmHg; *p* < 0.0001; n = 25) (Fig. 1c). This reduction was also apparent when comparing ICP eyes to fellow non-punctured control eyes at day 1 (13.21 ± 0.35 mmHg versus 17.69 ± 0.42; *p* < 0.0001). In ICP eyes compared to baseline, IOP was marginally but significantly reduced on day 3 (16.7 ± 0.79 mmHg; *p* < 0.05), normalizing by day 7. The difference in IOP between ICP and contralateral eyes was only observed on day 1. Intracameral injections of 2 µL PBS also showed reduced IOP at day 1 after injection (Supplementary Figure 1). At 24 hours following ICP, the corneas were fluorescein stained and puncture sites were found to be fully healed and without leakage. Immunostaining after 24 hours also showed full corneal wound healing and scar formation at puncture sites (Fig 1d).

### Outflow Facility and Aqueous Production are Unchanged After Intracameral Puncture

To assess the factors affecting aqueous humor dynamics at maximum IOP decrease (day 1), enucleated ICP and control eyes were subjected to *ex vivo* perfusions (n = 15 per group). Surprisingly, there was no difference in outflow facility between ICP and control eyes (2.15 ± 0.26 nl/min/mmHg versus 1.83 ± 0.18 ± 0.27 nl/min/mmHg; *p* = 0.31) (Fig. 2a).

**Figure 2:**
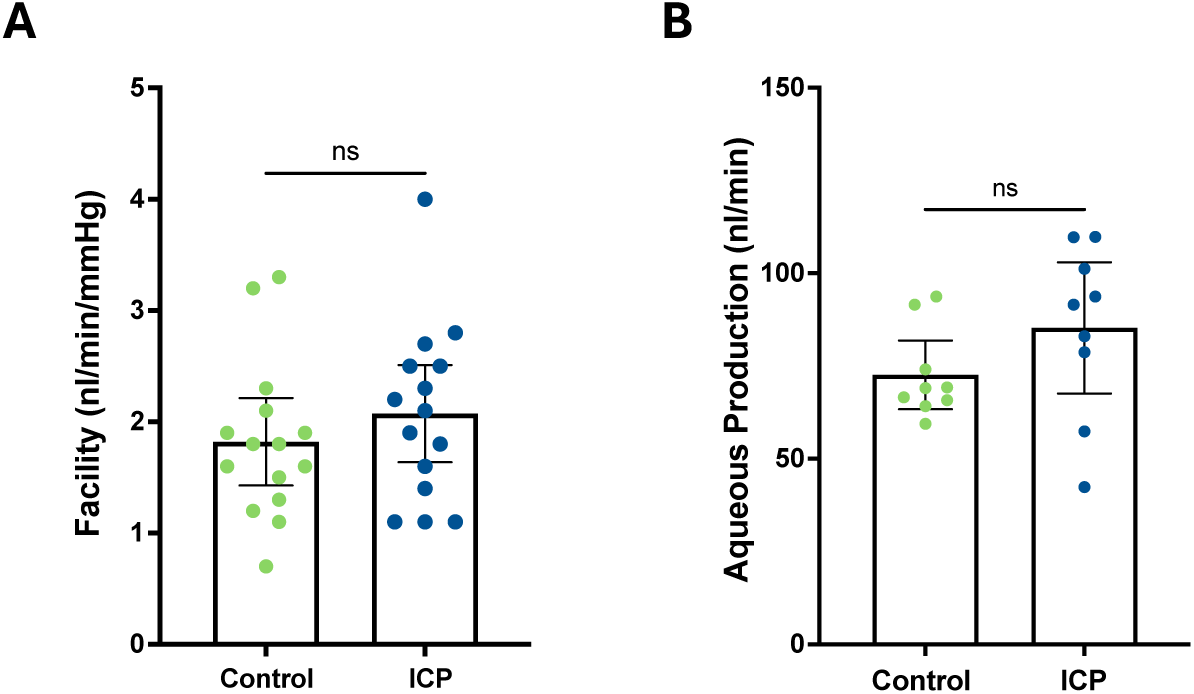
Outflow facility and aqueous humor production are unchanged at day 1 post ICP. (A) Outflow facility measured *ex vivo* using iPerfusion at day 1 following ICP in control and ICP eyes. (B) Aqueous production measured by fluorophotometry at day 1 in ICP eyes and contralateral controls. ICP: intracameral puncture. ns: not significant.

Next, aqueous humor production was measured on day 1. Using *in vivo* fluorophotometry, aqueous production rates were measured in ICP versus control eyes (n = 9 per group). Anterior chamber volume of 3 µL was used to calculate aqueous turnover.^17^ Results show that there was no statistically significant difference in aqueous production between ICP and control eyes (72.63 ± 4.01 nl/min versus 85.28 ± 7.66 nl/min; *p* = 0.16) (Fig. 2b). As a control, aqueous production was measured in dorzolamide-treated and control wild-type mice to validate reduced aqueous production by inhibiting carbonic anhydrase in ciliary body epithelium with dorzolamide (Supplementary Figure 2). We therefore conclude that ICP does not alter aqueous humor dynamics by means of outflow facility or aqueous production.

### Episcleral Vasculature Dilation After Intracameral Puncture

To investigate whether reduced IOP on day 1 after ICP could be attributed to changes in the distal portion of the conventional outflow tract, the diameter of episcleral vessels including distal venules (n = 125 measurements in 3 mice) (Fig. 3a) were measured. At day 1, there was an increase in the diameters of distal venules compared to contralateral control eyes (22.78 ± 0.53 µm versus 18.26 ± 0.47 µm; *p* < 0.0001) (Fig. 3b). This dilation of venules was consistently observed in all four quadrants of ICP anterior segment whole mount samples. Increased vessel diameter, measured at day 1, was also observed in the arterioles (15.64 ± 0.34 µm versus 13.7 ± 0.31µm; *p* < 0.0001) and capillaries (6.84 ± 0.13 µm versus 6.35 ± 0.13 µm; *p* < 0.01) of ICP eyes verses contralateral controls (Fig. 3b). These results indicate that ICP increases the caliber of the distal vasculature, likely reducing distal outflow resistance.

**Figure 3:**
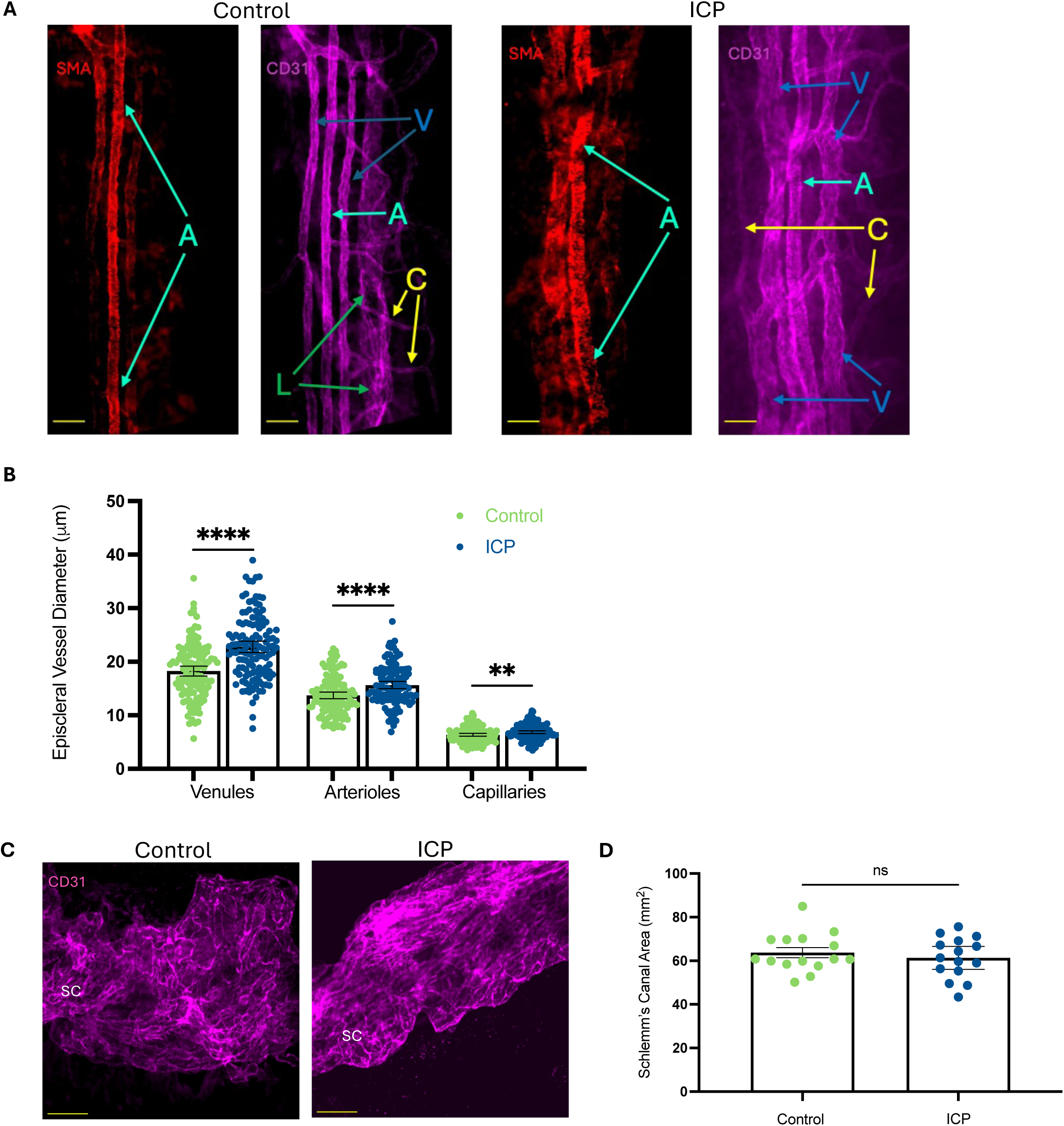
Episcleral vessels are significantly dilated while Schlemm’s canal is unaffected at day 1 post ICP. (A) Maximum projection *en face* confocal images of the episcleral vessels in control and contralateral ICP eyes at day 1. Arterioles are identified by striated anti-SMA staining. (B) Quantification of vessel diameters of episcleral venules, arterioles, and capillaries from tiled images in ICP and contralateral control eyes after 1 day. (C) Maximum projection images of Schlemm’s canal in control and contralateral ICP eyes after 1 day. (D) Schlemm’s canal area quantified in ICP eyes and contralateral controls. All scale bars = 50 µm. A (teal arrows): arterioles; C (yellow arrows): capillaries; L (green arrows): lymphatics; V (blue arrows): venules; SC: Schlemm’s canal; ICP = intracameral puncture. ** *p* < 0.003; **** *p* < 0.0001; ns: not significant.

Moving further upstream of aqueous humor drainage, the area of SC was also measured. Unlike the distal vessels, there was no difference in SC area between ICP and contralateral control eyes at day 1 (61.34 ± 2.45 µm versus 63.71 ± 2.28 µm; *p* = 0.48) (Fig. 3c, d). These results show no effect on SC area with ICP and corroborate prior results demonstrating outflow facility is unchanged.

### Macrophages are Recruited Following Intracameral Puncture

Macrophages are key sensors of tissue damage and immune surveillance.^13^ To determine if ICP affects macrophage quantity in the conventional outflow tract, macrophage numbers were counted in outflow tract tissues: TM, SC and DV, at baseline and 1, 3, 7 and 14 days after ICP (n = 15 per group). Macrophages in TM were localized in confocal sections with anti-SMA Ab staining, whereas macrophages in SC were localized with anti-CD31 Ab. DV macrophages adjacent to CD31^+^ vasculature were counted. Corresponding to the IOP reduction, there was an increase in macrophage density in the outflow tract beginning at 1 day following ICP and lasting up to 14 days. Specifically, Iba1^+^ macrophages were recruited to the episcleral and distal vasculature on days 1 (3077 ± 231 macrophages/mm^2^ versus 1802 ± 226 macrophages/mm^2^; *p* < 0.0005; n = 15) and 3 (3244 ± 175 macrophages/mm^2^ versus 2460 ± 253 macrophages/mm^2^; *p* < 0.05; n = 15), and returned to baseline values on day 7 (*p* = 0.25) in ICP versus control eyes (Fig. 4a, d).

**Figure 4:**
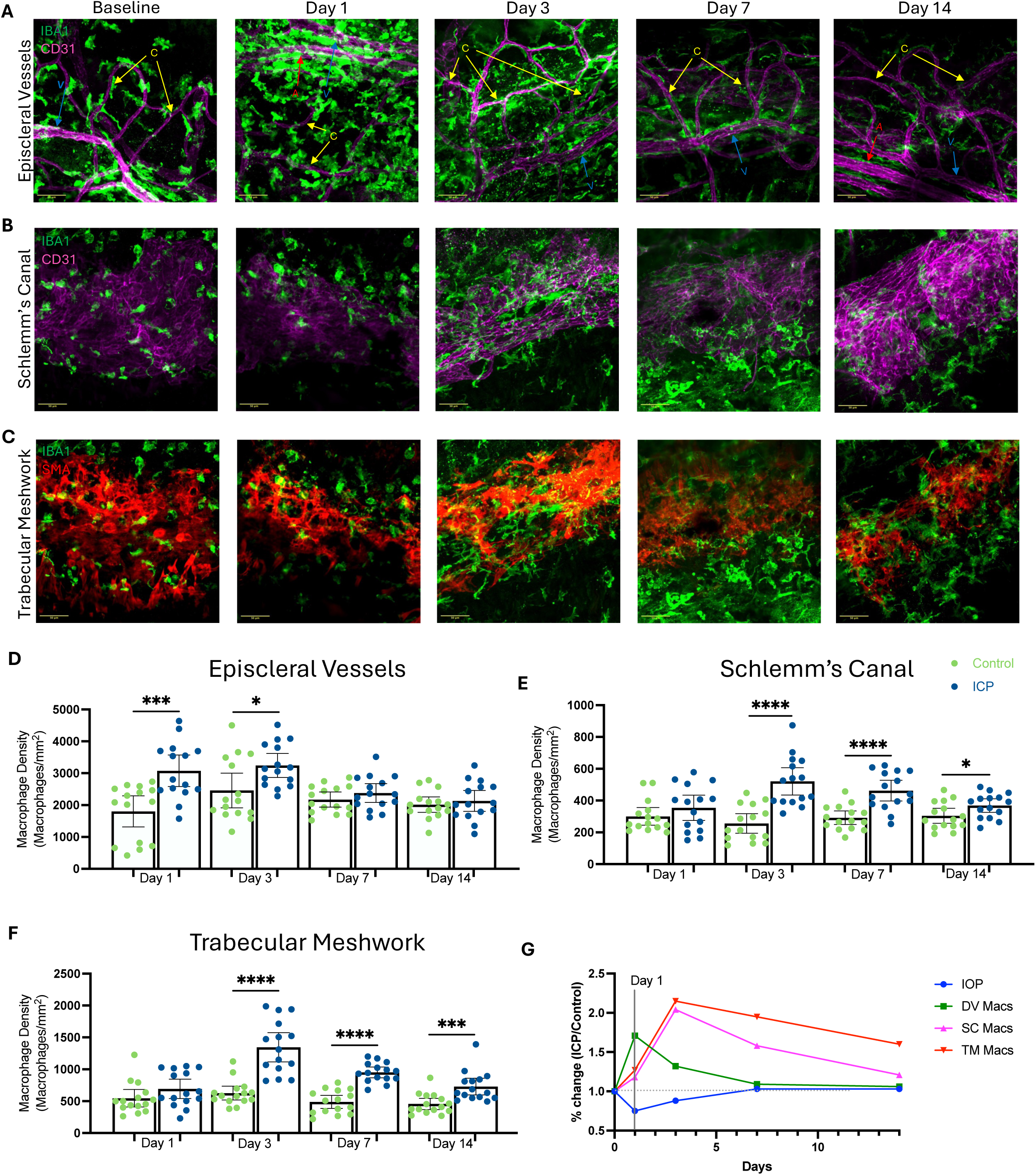
Macrophage density is significantly increased in outflow tract tissues following ICP. Maximum projection *en face* confocal images the episcleral vasculature (CD31^+^) (A), Schlemm’s canal (CD31^+^) (B), and trabecular meshwork (αSMA^+^) (C) with influx of macrophages (Iba1^+^) at baseline and days 1, 3, 7, and 14 post ICP. Quantification of macrophage density around DV (D), Schlemm’s canal (E), and in trabecular meshwork (F) at days 1, 3, 7, and 14 in ICP eyes and contralateral controls. (G) Overlay plot showing IOP versus macrophage spatiotemporal distribution after ICP (percent change in ICP macrophage density / control macrophage density at each time point). Scale bar = 50 µm. ICP: intracameral puncture; IOP: intraocular pressure; DV: distal vessels; SC: Schlemm’s canal; TM: trabecular meshwork; Macs: macrophages. * *p* < 0.04; *** *p* < 0.0002; **** *p* < 0.0001.

Macrophages significantly infiltrated Schlemm’s canal 3 days following ICP versus contralateral controls (520 ± 40 macrophages/mm^2^ versus 256 ± 29 macrophages/mm^2^; *p* < 0.0001; n = 15) and this increase persisted for up to 14 days (*p* < 0.05) (Fig. 4b, e). Finally, ICP eyes showed a significant increase in macrophages within the TM at 3 days (1346 ± 107 macrophages/mm^2^) and lasting up to 14 days following ICP (729 ± 61 macrophages/mm^2^), compared to contralateral controls (626 ± 50 macrophages/mm^2^ and 456 ± 41 macrophages/mm^2^, respectively; *p* < 0.001 at both timepoints; n = 15) (Fig. 4c, f).

A comparison of IOP and spatiotemporal dynamics of macrophages in the outflow tract showed that the reduction in IOP at day 1 corresponded with recruitment of macrophages to episcleral vessels (Fig. 4g). Macrophage recruitment to the ciliary epithelium 1 day after ICP was also analyzed in cross-sections of ciliary body (4 mice; Supplementary Figure 3a). Results show that there was no difference in the number of macrophages per unit area within the ciliary body 1 day after ICP (1355 ± 105 macrophages/mm^2^ versus 1211 ± 113 macrophages/mm^2^; *p* = 0.35; n = 20 sections, 4 mice per group) (Supplementary Figure 3b).

## Discussion

Intracameral injections are a commonly used technique to deliver drugs, viruses and other biomedical agents to the anterior chamber and outflow tract tissues. However, the effect of intracameral injections on IOP and aqueous humor dynamics has not been studied. In the present study, we observed that intracameral injections lead to a temporary reduction in IOP, a simultaneous recruitment of macrophages to the distal portion of conventional outflow tract tissues that corresponds with dilation of distal vessels, but no change in outflow facility or aqueous production. While IOP and macrophage abundance return to baseline levels by day 7 in the distal portion of the conventional outflow tract, macrophage numbers remain elevated in TM and SC two weeks after ICP.

Following ICP, IOP was significantly decreased at day 1 (24 hours), marginally reduced at day 3 and returned to baseline values from day 7 onward. This effect on IOP was not due to leakage of aqueous humor as corneal puncture wounds were completely healed at day 1. As a small volume of aqueous humor refluxes into the empty glass needle, reduced IOP was also not due to the loss aqueous from the ICP itself. The entire volume of aqueous humor in the anterior chamber (approximately 3 µL) turns over in ∼50 minutes, given that aqueous is produced at a rate of ∼60 nl/min.^17^ Thus, the reduction in IOP is due to alteration in aqueous humor dynamics, which has not been previously evaluated following ICP.

We first assessed outflow facility and found no change at day 1 after ICP. This finding was consistent with Li *et al*. who showed that intracameral injections of adenovirus encoding GFP did not affect outflow facility 3-6 days following injections.^1^ While the effect of intracameral injections on outflow facility has not been specifically studied in humans, Wen *et al*. demonstrated that intravitreal injections of anti-vascular endothelial growth factor (anti-VEGF) did not influence outflow facility in patients receiving <10 injections.^19^ We next determined whether ICP affects aqueous production. Reduced ciliary body function (i.e., ciliary body shutdown) has been proposed as a mechanism for reduced IOP, particularly in cases of inflammation.^20^ However, we found that there was no change in the rate of aqueous production at day 1 after ICP.

Given that outflow facility and aqueous production were unchanged, we next examined the morphology of the episcleral vessels, as currently there is no established method to measure distal outflow resistance or a reliable method to measure episcleral venous pressure in pigmented mice. Our results showed a significant increase in the caliber of episcleral vasculature (i.e., venules, arterioles and capillaries) on day 1 after ICP. The distal outflow tract is comprised of the collector channels, intrascleral vessels and episcleral vessels. Importantly, the distal outflow tract can account for 25-50% of outflow resistance.^21^ Pharmacologic vasodilation of distal vessels with nitric oxide (diethylenetriamine-NO) increases distal outflow facility in post-trabeculotomy human and porcine anterior segments.^22^ Thus, our results suggest the ICP may be acting at the level of the distal vasculature to affect IOP.

Macrophages are key immune sensors after tissue injury. Therefore, we evaluated the spatiotemporal dynamics of macrophages in the outflow tract tissues following ICP. We found macrophages are recruited to the outflow tract tissues with a significant increase in macrophage density around episcleral vessels at days 1 and 3 with a return to baseline at later time points.

There was a subsequent increase in SC and TM macrophages at days 3, 7 and 14. We observed an increase in macrophages around episcleral vessels concurrent with reduced IOP at day 1 (Fig. 1g). While macrophages reside in the outflow tract tissues in homeostasis,^10,23^ our results suggest, following ICP, monocytes are recruited first to the episcleral vessels, which have direct access from the venous circulation. These leukocytes then extravasate from the episcleral venules where they differentiate into macrophages, and then with time, travel to the SC and TM. It has recently been discovered that macrophages directly affect IOP, as depletion of resident macrophages alters IOP homeostasis.^23^ Furthermore, drugs that modify the immune response (i.e., prostaglandin analogs and corticosteroids) also affect IOP.^24–26^ We speculate that macrophages at the level of the distal vasculature could potentially affect distal outflow by directly interacting with the vascular smooth muscle cells and/or surrounding extracellular matrix,^27^ or macrophages surrounding the distal vasculature could be secreting cytokines to affect vascular tone.^28,29^

Limitations of this study include the investigation of intracameral delivery only and thus our results cannot be extended to intravitreal injections. We studied macrophages as they have been shown to be important in IOP homeostasis,^23^ but the response and significance of other immune cell types have not been elucidated. Furthermore, we did not study macrophages by ontogeny in order to confirm that recruited macrophages are monocyte derived. Finally, future functional studies are needed to determine whether monocyte recruitment directly affects the IOP response on day 1.

In this study, we show that IOP is reduced within 24 hours after intracameral puncture, while outflow facility and aqueous production are unchanged. This reduced IOP is accompanied by macrophage influx into the outflow tract tissues. At day 1, macrophages are recruited to episcleral vessels with concurrent dilation of episcleral vessel caliber, suggesting short-term alterations in IOP after ICP may be due to the immune response at the level of the distal vasculature. Taken together, our study demonstrates the importance of considering the method of drug delivery to the eye, particularly when analyzing conventional outflow homeostasis and the immune response.

## Acknowledgements

The authors thank Bruce Ishimoto, Carol Toris and Shan Fan for their assistance with fluorophotometry.

## Supplemental Figure Legend

**Supplementary Figure 1:**
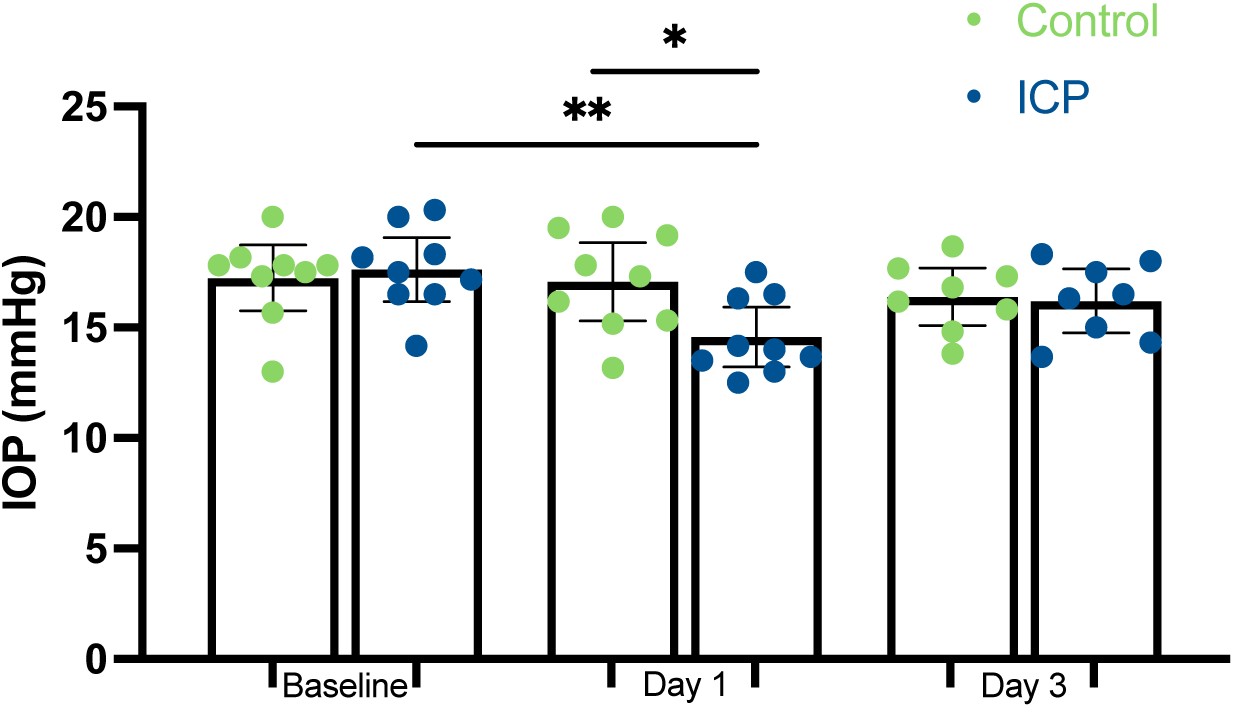
Intracameral injection reduces IOP at day 1. IOP in C57BL/6 mice, age 3-5 months, at baseline and days 1 and 3 following intracameral injection of 2 µL PBS. * *p* < 0.04; ** *p* < 0.003.

**Supplementary Figure 2:**
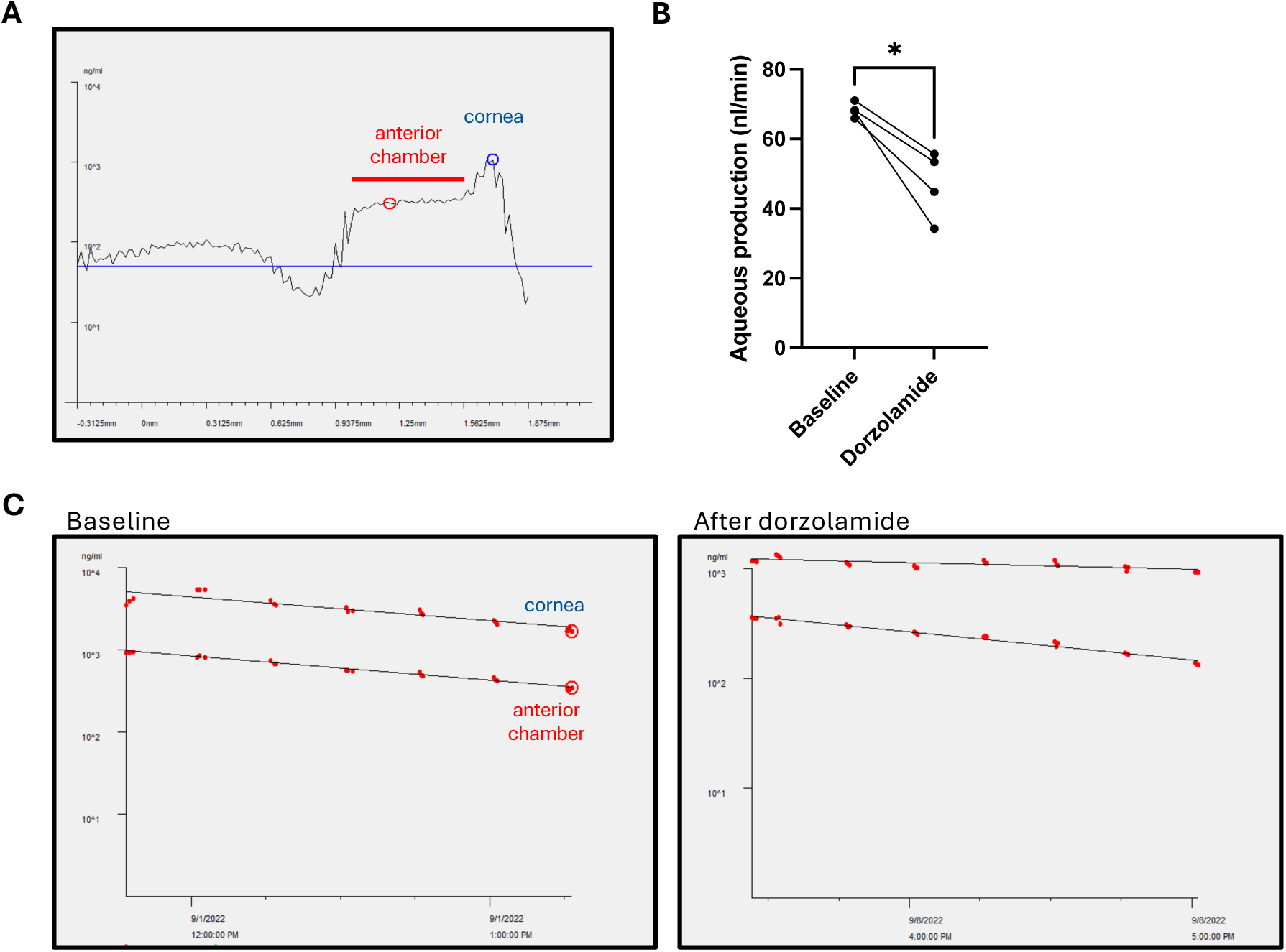
Aqueous production is reduced with dorzolamide treatment. (A) Representative line scan of single measurement from an adult wild type C57Bl/6 mouse using the Fluorotron^TM^ Master Fluorophotometer (Ocumetrics, Mountain View, CA). (B) Aqueous production in C57Bl/6 wild type mice (n = 3) at baseline and after dorzolamide treatment. **(C)** Collated fluorophotometry measurements in a wild type mouse at baseline and after dorzolamide eye drops. * *p* < 0.04.

**Supplementary Figure 3:**
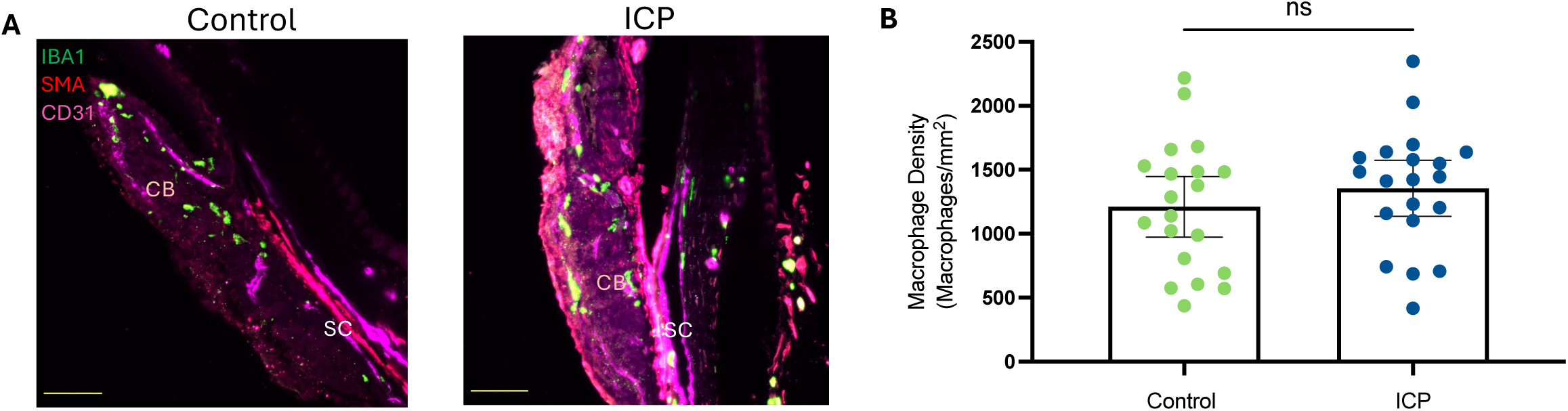
Macrophage density is unchanged in the ciliary body after ICP. (A) Immunofluorescence cross sections of ciliary body with Iba1^+^ macrophages at baseline and day 1 following ICP. (B) Quantification of macrophage density in ciliary body at day 1 in ICP eyes and contralateral controls. Scale bar = 50 µm. CB: ciliary body; SC: Schlemm’s canal; ICP: intracameral puncture. ns: not significant.

